# Comparing Cell Division- and Cell Reproduction-based Cell Lineage Analysis for Early Embryogenesis of *Caenorhabditis elegans*

**DOI:** 10.1101/199604

**Authors:** Shi V. Liu

## Abstract

Cell lineage analysis holds important stakes for understanding heredity and cell differentiation. Conventional cell lineages are reconstructed according to a cell division doctrine of one mother cell dividing into two daughter cells. An alternative cell lineage reconstruction method followed a cell reproduction discovery of multiple daughter cells reproduced from a same mother cell. To see which reconstruction method reflects reality of early embryogenesis of *Caenorhabditis elegans*, a side-by-side comparison was made between two methods. Here I show cell division-based lineage distorted reality and failed in revealing any true genealogy. Cell reproduction – based lineage conformed to reality with exact same number of cells in every developmental stage under examination and showed clear genealogical relationship. A paradigm-shift from cell division-to cell reproduction-based cell lineage analysis is necessary for correct understanding of developmental biology and will lead to a revolution in cell biology and life science.

## INTRODUCTION

Cell lineage analysis (CLA) holds important stakes for understanding total heredity comprising inheritance of genetic as well as epigenetic information. A faithful CLA should not only fit with the reality in the number of cells under examination but also truthfully reflect genealogical relationship among those cells. Unfortunately, most CLAs have failed in these aspects. Conventional CLAs following a cell division (CD) doctrine of one mother cell dividing into two daughter cells even excludes *bona fide* genealogical linkage in cell lineage reconstruction (Sulston and Horvitz, 1977; Sulston et al., 1983; Sulston, 2003). In contrast, novel CLAs based on a discovery that a mother cell can reproduce more than once to produce different daughter cells have enabled revelation of genealogy in cell lineages (Liu, 1999 and 2004a, b, Liu and Zhang, 2004a, b, Liu, 2005a, b and Liu, 2011a, b). Unfortunately, these CR-CLAs have been neglected and the mainstream is still dominated by the CD-based CLAs (CD-CLAs) (Liu, 2012a).

In order to break through a stone-walled CD-restriction on biological research and harvest tremendous value of CR-enlightened life science for biomedicine it is necessary to perform some side-by-side comparisons between CD-based and CR-based understanding of cell life. Directly contrasting CLAs from the CD-and the CR-perspective may accelerate the recognition of the deficiency and even delusion of the CD dogma and thus promote the acceptance of an insightful and truthful life science based on CR discoveries.

This direct comparison may be effectively made in a model organism with which rich information is available on its developmental biology. In addition, this comparison may be best performed in a forward rather than backward CLA. Ideally this forward CLA should be started from the very beginning of a multicellular life.

The invariant cell composition of *Caenorhabditis elegans* (*C. elegans*) (Sulston et al., 1983) and the availability of even some critical information (Zacharias and Murray, 2016) for making reliable mother-daughter (M-D) assignment made the beginning stage of *C. elegans* embryogenesis a suitable object for performing this forward CLA. This report presents some key results that directly compare CD-CLA with CR-CLA on cell lineage reconstruction for the first 100 minutes of the *Ce* embryogenesis and, more importantly, discuss these results with connections to previous publications criticizing on CD dogma and promoting CR insights. Because the same sets of data, obtained from respectful publications, were used for various comparisons between CD-CLAs and CR-CLAs, the comparisons were based on an objective basis. Thus, the pros and cons of contrasting CLAs should be self-evident.

## RESULTS

### Bifurcating Dendrogram Reconstructions

Figure 1A shows a typical CD-based bifurcating dendrogram for cell lineage reconstructed by condensing the top six levels in the master dendrogram showing complete embryogenic cell lineage of *Ce* (Sulston *et al*., 1983). There are 30 cells identified in the sixth level of this condensed dendrogram, rather than 32 cells as usually expected from synchronous cell divisions. Figure 1B shows an unconventional CR-based bifurcating dendrogram for these same 30 cells named according to the CR scheme [Liu, 1999] described in the Methods. In performing the CR-CLA an assumption was made that all the P_n_ cells depicted in the CD-CLA were the same mother cell captured at the different developmental stage after n^th^ reproduction. For assignment of M-D in other division pairs, an assumption of “left mother, right daughter” was made. But these M-D assignments may be totally wrong as shown in later CR-CLAs.

**Figure 1.**
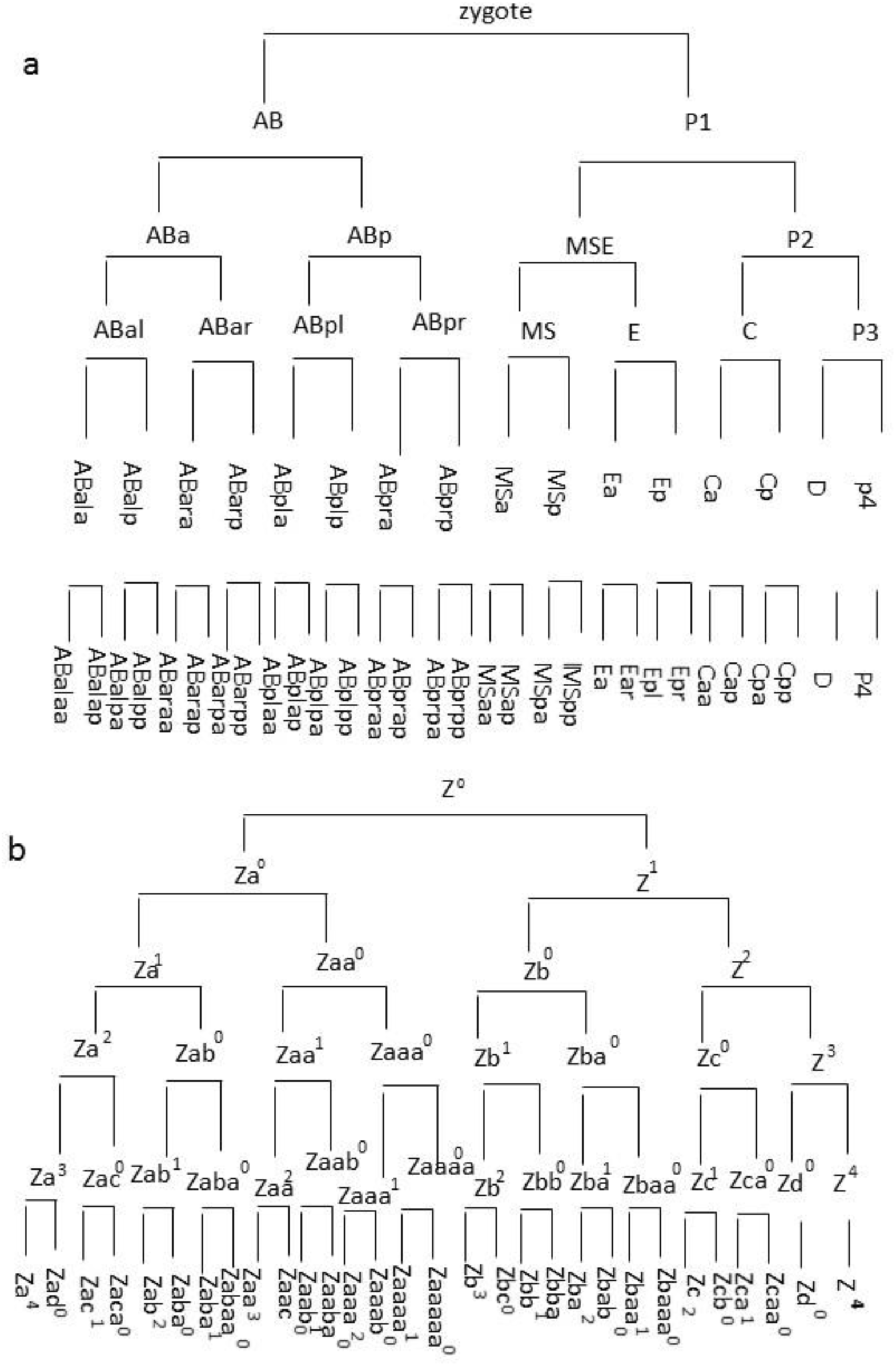
Dendrogram presentation of cell lineage for early stage embryogenesis of *Caenorhabditis elegans* according to cell division (A) and cell reproduction (B) principles, respectively.

An evident difference between the CD- and the CR-based cell lineage dendrogram is that CD-CLA dendrogram treats all division-paired cells as “sisters” and all cells on the same dendrogram level as “descendants” within a same “generation” (Figure 1A). However, this is not the case in the CR-based dendrogram in which every reproduction-pair bears an M-D relationship and cells at a same dendrogram level comprise an ancestor mother cell and its descendant(s) which can spread over multiple generations (Figure 1B).

### Pedigree Tree Reconstructions

The above distinctions between CD-CLA and CR-CLA became even clear when cell lineages were depicted as a pedigree tree. Figure 2A shows a pedigree tree constructed for 30 cells uniquely identified in the 6^th^ horizontal level of the above CR-CLA dendrogram (Figure 1B). Compared with the dendrogram presentation of cell lineage, pedigree representation of cell lineage shows even clearer genealogical relationship among all cells under the examination. There were a total of 6 generations among these 30 cells and their generation distribution is shown in the insert table in Figure 2A.

**Figure 2.**
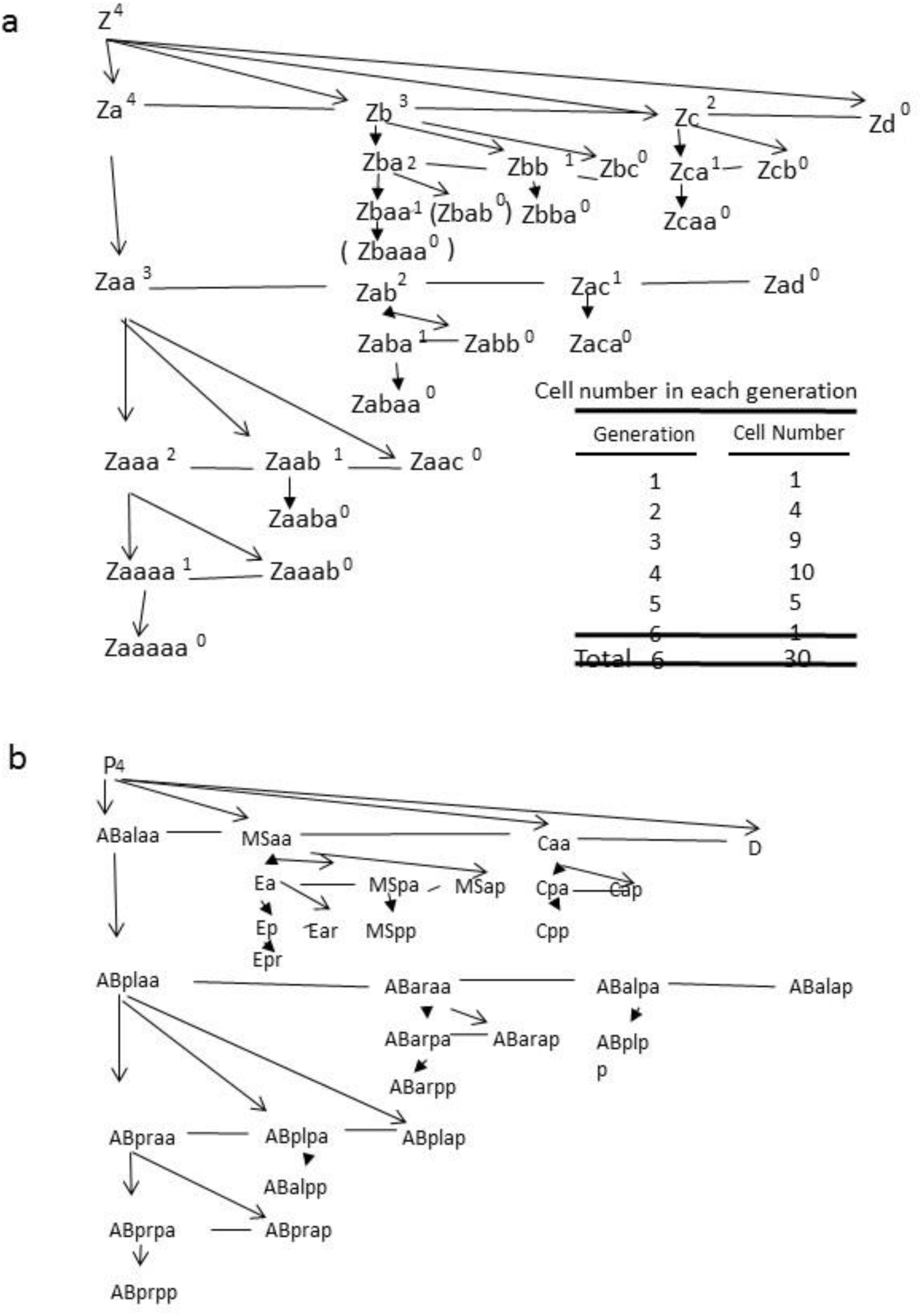
Pedigree trees of cells named according to cell reproduction (A) and cell division (B) principles for the early stage embryogenesis of *Caenorhabditis elegans*. Arrowed lines indicating cell reproduction while horizontal lines linking sibling cells in the same generation.

Because CD-CLA itself could not generate a pedigree tree CD-named cells occupying equivalent positions between CD-CLA and CR-CLA dendrogram were forced onto the CR-CLA-derived pedigree tree (Figure 2B). However, this forced pedigree tree does not make any genealogical sense by the CD-based cell names.

The above contrast demonstrates that, while CR-CLA can easily shed rich genealogical information, CD-CLA does not have an intrinsic capability of revealing any true genealogy.

### 3D Spatial Reconstructions

The deficiency of CD-CLA became even more serious when its results were put into a reality check. The landmark publication of embryonic cell lineage for *C. elegans* contains a diagram showing the spatial locations of 28 cells (Sulston et al., 1983), rather than 30 cells as would be collected from the above dendrogram cell lineage reconstruction. Later studies confirmed that, within the first 100 minutes of *Ce* embryogenesis, there are indeed just 28 cells derived from zygote and there is no death/loss of cell over the course (Heid, et al., 2002; Giurumescu and Chisholm, 2011). Thus, if a CLA really works, it should be capable of directly identifying all genealogical relationship among these cells. However, when CD-CLA was applied to a 3D spatial reconstruction of cell lineage, the only genealogical relationship that could be identified was a “sister” relationship (Figure S1A) which depends on the correctness of the assumption that division-paired cells are “twin daughters”. However, this assumption is wrong when CR-CLA actually identified these “sister” pairs as M-D pairs (Figure S1B).

Unlike CD-based CLA which would treat all 28 cells as “descendants” of one same “generation”, CR-CLA showed that these 28 cells belong to 6 different generations (Figure 2A) with Z^4^ as the oldest cell which had reproduced 4 times to give birth of Za, Zb, Zc, and Zd (Figure S1B). The youngest cell is Zaaaaa ^0^ which belongs to the 6^th^ generation and has not yet experienced any cell reproduction. However, it should be pointed out that, due to a potential mistake in the M-D assignment which will be seen in later analysis, the placement of some descendant cells in this 3D cell lineage reconstruction might be still be wrong.

### 4D Spatiotemporal Reconstructions

Detailed time record is very useful for cell lineage reconstruction. A 4D spatiotemporal dynamics of cell lineage formation was thus reconstructed from time-stamped images of the very early stage of *C. elegans* embryogenesis [Verbrugghe and Chan, 2011]. The sequential formation of 1-, 2-, 3-, and 4-cells were labelled according to the CD-(Figure S2A) and CR-(Figure S2B) cell naming rules, respectively. However, it is worth of warning that, even with this 4D CLA, the M-D assignment for the Za-Zaa pair in the CR-CLA is still uncertain because critical cell age information was lacking for certifying which one is the real mother and which one is the real daughter.

### CR-based “5D” Reconstruction

Since it is hard to find cell age information in conventional CD-CLAs, other information such as cell polarity memory, cell contact history, gene expression profile, and signaling pathway may be used for making some “educated” guess on the M-D identification. Collectively this set of information was regarded in this study as the 5^th^ dimension in cell lineage reconstruction for approaching genealogical reflection. A recent review entitled “combinatorial decoding of the invariant C. elegans embryonic lineage in space and time (Zacharias and Murray, 2016) contained some such 5^th^ dimension information. While this review still presented a CD-based CLA for the cell lineage formation from zygote to the 16-cell embryonic stage (Figure 3A), a CR-based cell lineage reconstruction was created by taking the relevant 5^th^ dimension information into consideration of M-D assignment for all division-pairs (Figure 3B).

**Figure 3.**
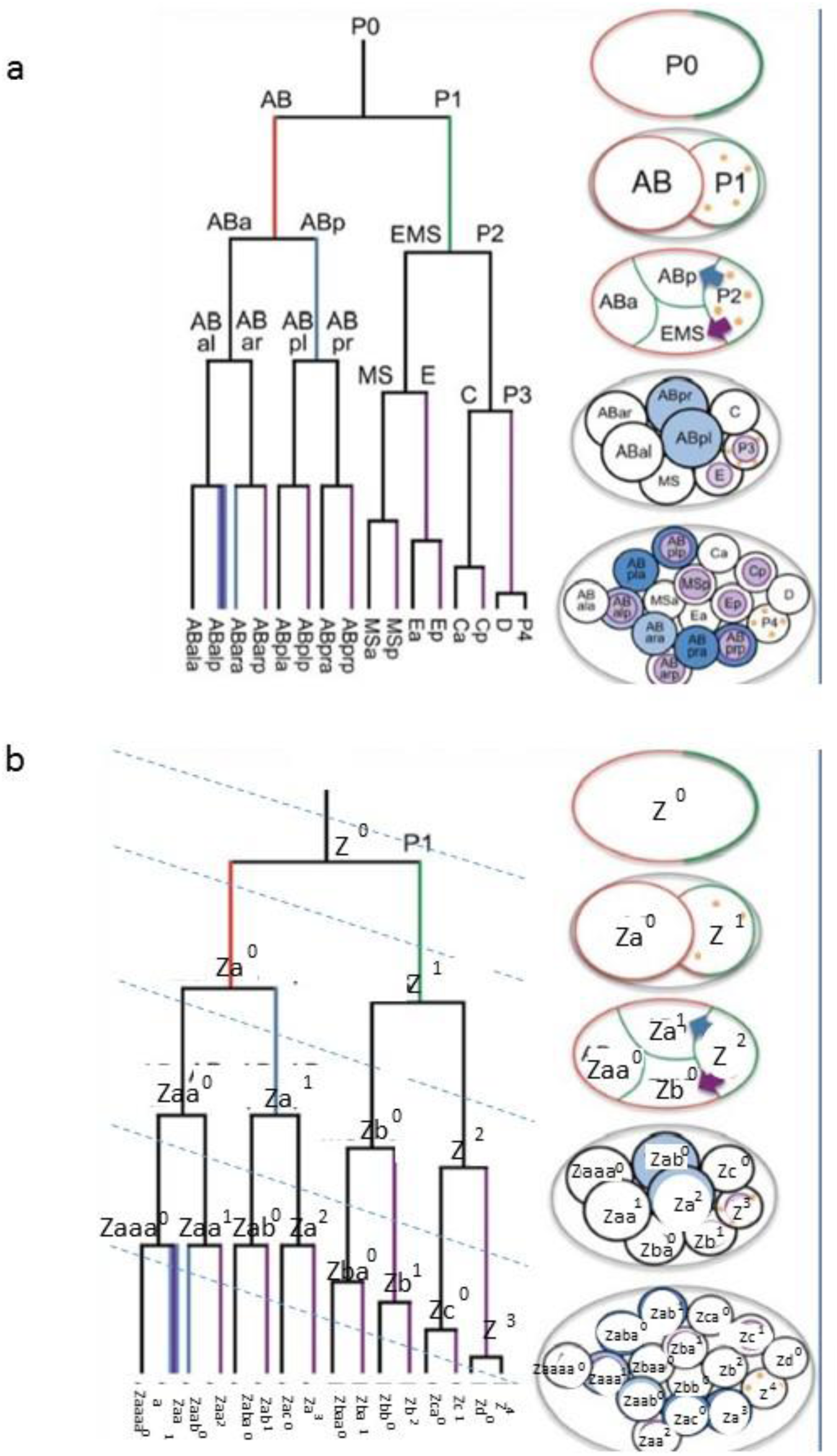
5D spatiotemporal dynamics of cell lineage during 1-to 16-cell stages of *Caenorhabditis elegans* embryogenesis with cells named according to CD-(A) and CR-(B) scheme.

In comparing this 5D reconstruction of the 16-cell cell lineage (Figure 3B) with the previous 3D reconstruction of the 28-cell cell lineage (Figure S1), it was noticed that the 5D reconstruction showed some consistent spatial pattern in the distribution of generation-different cells (Figure 3B), rather than a more random distribution in the previous 3D reconstruction (Figure S1). In this 5D reconstruction, all daughters of Z (Za, Zb, Zc, and Zd) were found in close distance to their mother cell. The daughter cells of Zx were generally found in proximity to their respective mother cell. Thus, a kind of generation progression was reflected in the spatial arrangement of the embryonic cells. An overall “cell age” polarity was seen in the 16-cell embryo: the oldest cell (Z^4^) and the youngest cell (Zaaaa^0^) were found at the opposite ends of the embryo (Figure 3B).

This 5D reconstruction of cell lineage also revealed a common feature: a mother cell was usually found at the posterior side and a daughter cell was usually found at the anterior side of a given M-D pair (Figure 3B). Thus, the “left mother, right daughter” assumption made for previous CR-CLAs was apparently wrong and thus should be corrected as “right mother, left daughter”. When this “right mother, left daughter” rule was used for renaming cells in the CD dendrogram (Figure 3A), the CR dendrogram showed the placement of the cells matching those in the spatial depiction (Figure 3B), with the oldest and the youngest cell on the two sides of the dendrogram.

Also noticed in the CR-dendrogram was some regularity in the timing of the reproduction waves, indicated by the dashed lines cutting across the CR dendrogram (Figure 3B). A subsequent reproduction of the mother zygote cell was always later than the reproduction of its previous daughter and this lag in the reproduction timing was smaller for descendant cells of Za than for the descendants of Zb (Figure 3B). This reproduction asynchrony across the same level of dendrogram could explain why there were only 30 cells shown in the previous CLAs (Figure 1 and 2) and only 28 cells were in the “32-cell” stage embryo (Figure S1).

### Cell Lineage and Developmental Topology

From above analyses, it became clear that basic principles of CR-CLA could be summarized into a simple diagram (Figure 4A). In this diagram, the so-called “generation” as mistakenly believed in CD-CLA is correctly identified as developmental stage. The so-called “division” asynchrony is perceived as timing differences in reproduction among different cells. Cells captured in a same reproduction wave can be different in genealogical generation and close in chronological age.

**Figure 4.**
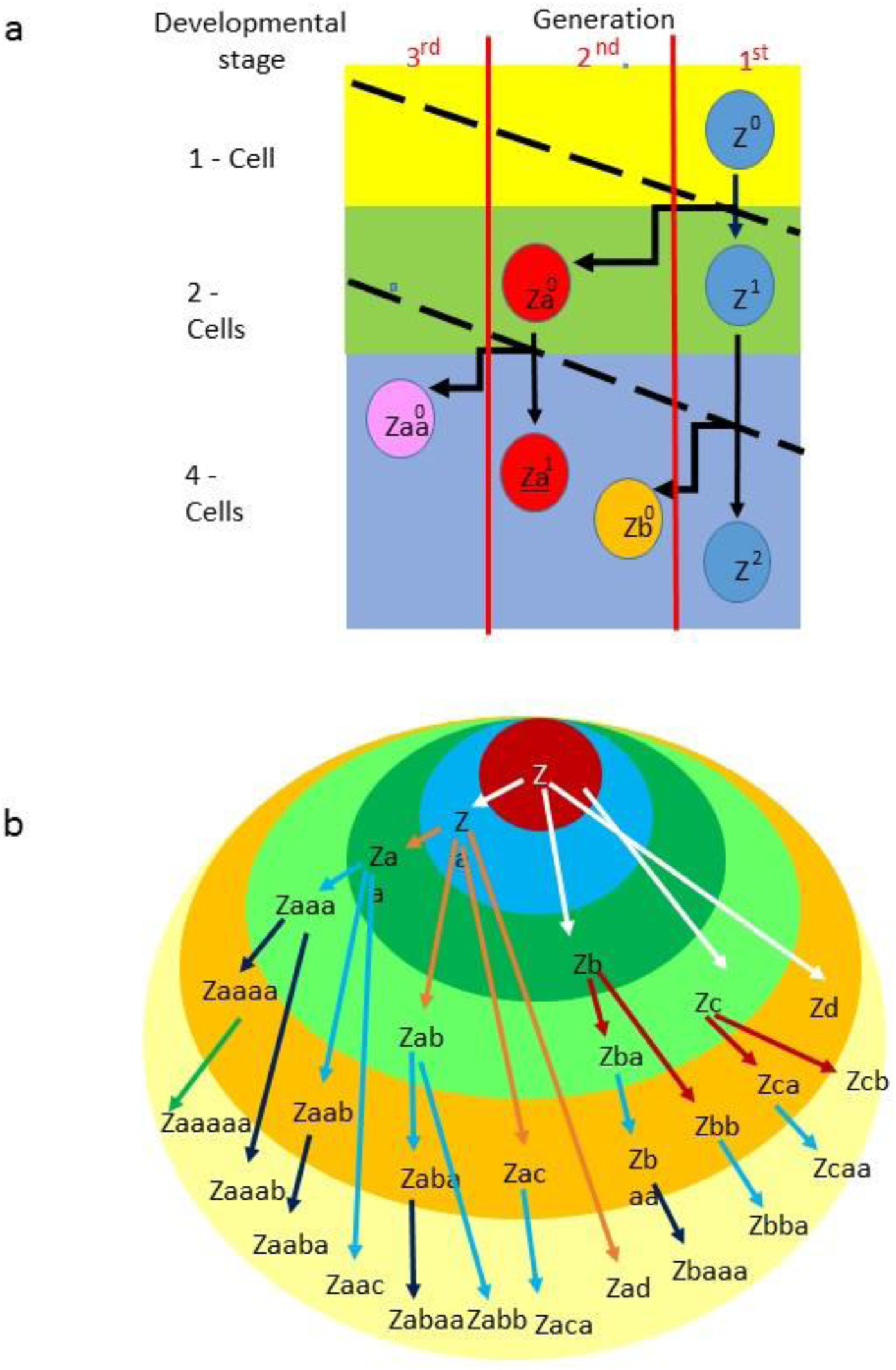
A schematic summary of the core features of CR-CLA (A) and a diagram showing developmental topology (B) of a 28-cell embryo of *Caenorhabditis elegans*. In the schematic summary (A) the different development stages were shown as blocks in different color and the different generations of the cells were separated with vertical lines. Cell reproduction link was indicated with zigzag arrow. Cell transition from an earlier to a later stage was indicated by vertical arrow. The dash lines separate different waves of cell reproduction. In the topology diagram (B), different waves of cell reproduction were shown as isochrones with different colors which also represent different chronological ages. Cell lineages were shown by cells linked with arrowed lines in different colors as white, red, blue, black, and green to represent linkage up to the first, second, third, fourth, and fifth generation, respectively.

Furthermore, some regularity was detected in the developmental topology enabled with the CR-CLA (Figure 4B). A multicellular organism develops from a single cell by consecutive cell reproduction waves, shown as spatially expanding rings (isochrones) over increasing time outwards. New and thus younger cells are added into the multicellular organism so the topological space of a developing multicellular organism increases. Cells collected in each isochrone have similar chronological ages even though they may belong to different generations due to the reproduction asynchrony among different cell lineages. This cell reproduction asynchrony is also an indication of cell differentiation besides other conventionally recognized aspects of cell differentiation.

### Cell Age Heterogeneity in Early Embryo

With the above CR-CLA principles in mind and recognizing a potential systematic error in M-D assignment in the earlier CR-CLAs the dendrogram for the 28-cell *C. elegans* embryo was updated with a consistent M-D assignment according to the “right mother, left daughter” assumption (Figure 5A). This change did not affect pedigree structure because its genealogical revelation of cell lineage is not affected by cell placement. However, this correction in M-D assignment had a dramatic impact on the 3-D cell lineage mapping (Figure 5B). By correcting cell names according to the updated dendrogram, a very regular placement of cells showing M-D proximity and generation gap was seen (Figure 5B), in contrast to a more random distribution of cell lacking such regularities (Figure S1B). The same “cell age” polarity as seen for the above CR-CLA on the 16-cell embryo (Figure 3B) was consistently shown here for the 28-cell embryo (Figure 5B) with the oldest (Z^4^) and youngest cell (Zaaaaa^0^) on the opposite poles.

**Figure 5.**
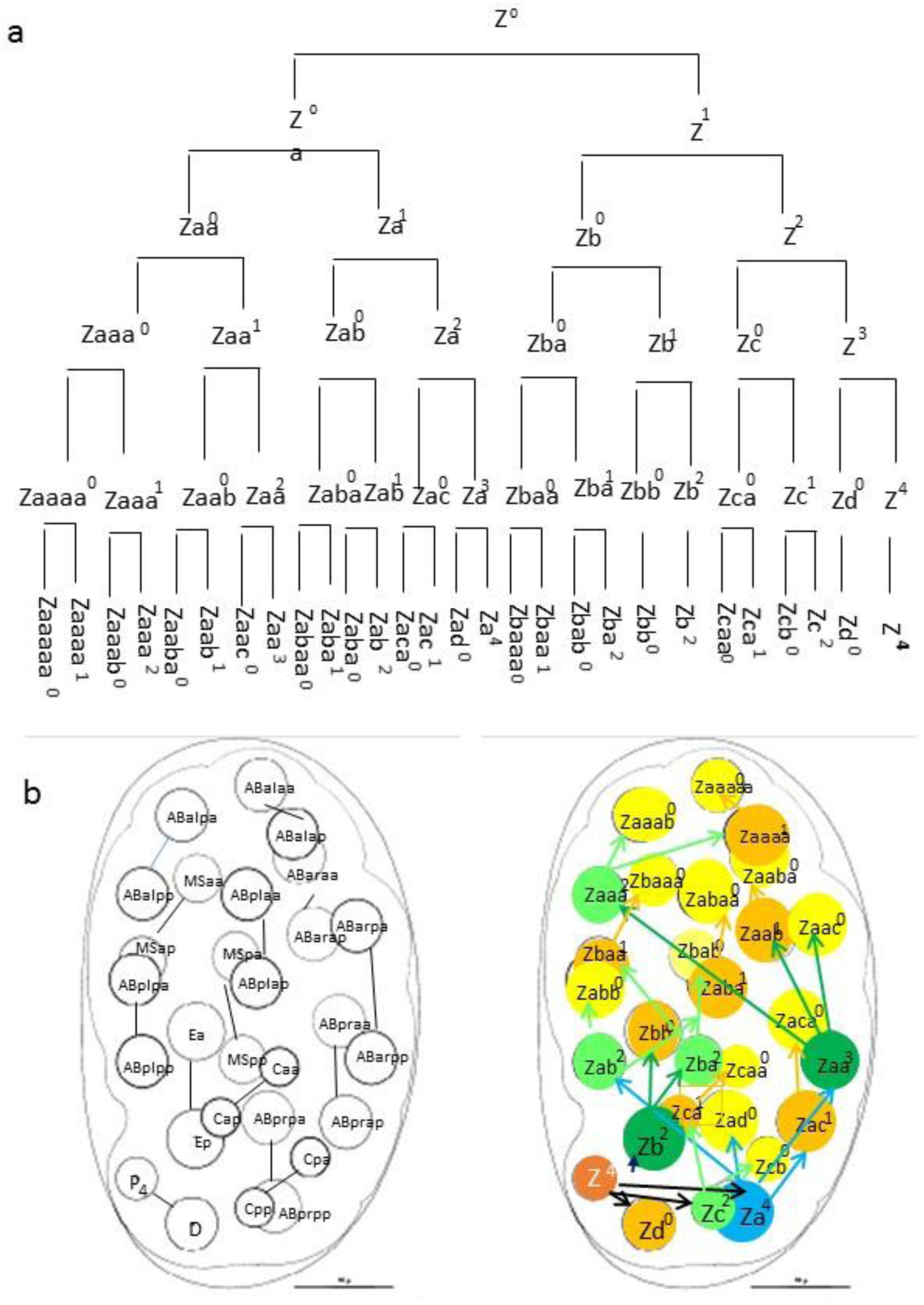
CR-based lineage dendrogram (A) and 3 D map (B) for the 28-cell embryos by CD-(left) and CR-(right) principles, respectively. The colors of cells in the CR-CLA map were based on the same set of colors used for showing developmental topology (Figure 4B).

As a matter of fact, when the same series of age-meaning colors used for depicting the sequential waves of cell reproductions in the developmental topology (Figure 4B) was used for filling cells in the 3D genealogical mapping, the cell age asymmetry in the 28-cell embryo became more obvious (Figure 5B). The germline cell pole contains more early-generation and chronologically older cells while the non-germline cell pole contains more later-generation and chronologically younger cells.

### Cell Differentiation from CD-and CR-Perspective

A very important feature in *Ce* embryogenesis is its very early cell differentiation which resulted in five different “founder cells” (Sulston, 1983) (Figure 6A). These different founder cells were regarded as ancestors of different sub-lineages which contribute to the formation of different tissues. From a CD perspective, these cell differentiations are achieved through consecutive cell-fate-change asymmetric cell divisions which result in two different daughter cells at each division. Thus, in the CD-CLA, different P cells are not only differentiated from their respective “sister” cells but also differentiated in themselves over time (Figure 6A). However, in the CR-CLA, all P cells were identified as the same mother cell having experienced sequential rounds of cell reproduction (Figure 6B). So there is no real cell-fate-change differentiation in this mother zygote cell even though its status might have changed in some other ways such as by normal aging and by abnormal mutation. On the other hand, the CR-CLA indicates that differentiating cell reproduction likely happens in different ways in different rounds of cell reproduction of zygote to form different daughter cells that serve as distinctive “founder” or stem cells leading to different sub-lineages contributing to different tissues (Figure 6B).

**Figure 6.**
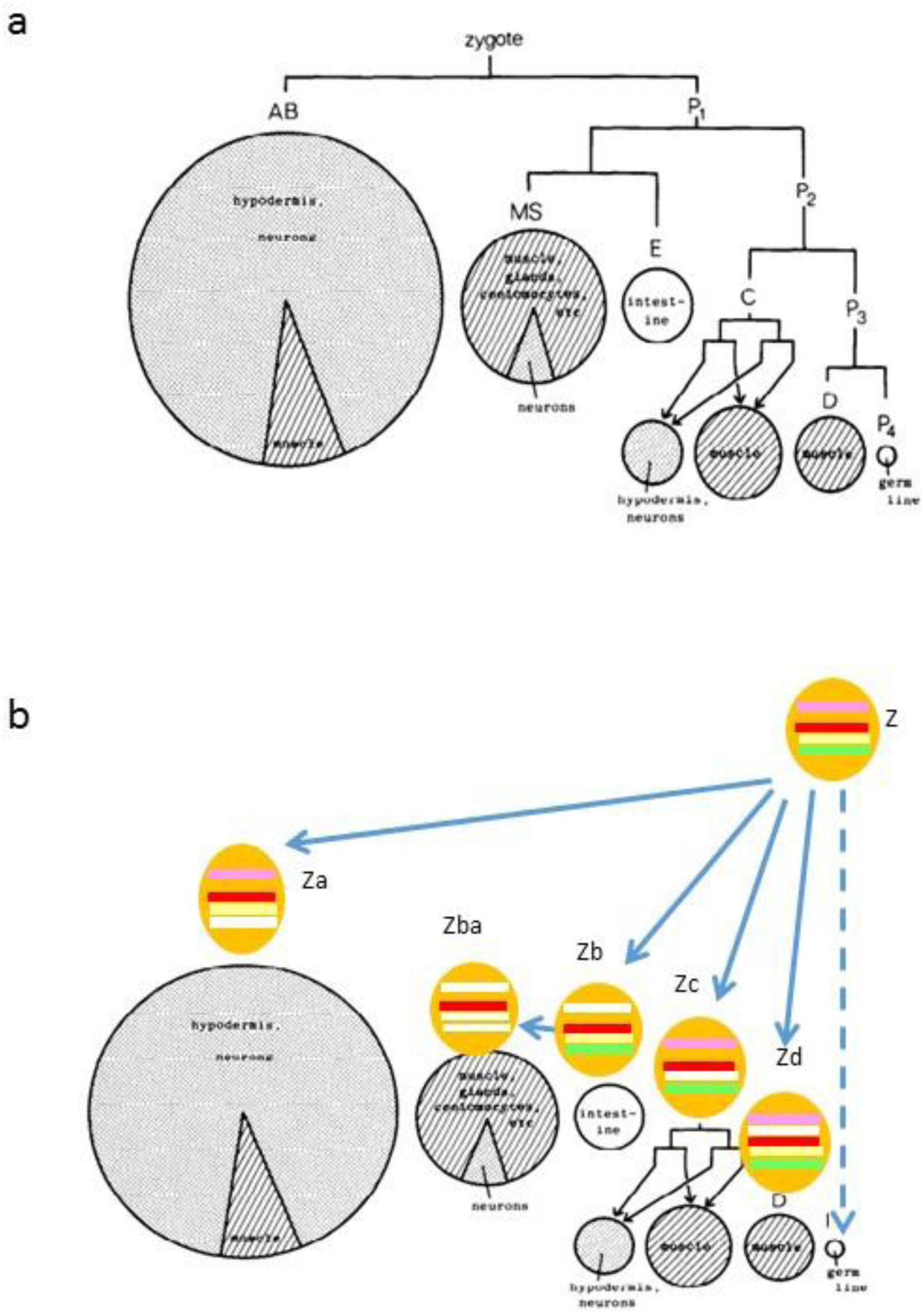
Cell differentiation and tissue formation from CD-(A) and CR-(B) based cell lineage formation. In the CR-cell lineage representation, the totipotency of zygote (Z) was shown with a full set of color bars. Upon differentiating cell reproduction (indicated by arrowed line linking reproduction pairs), different pluripotent stem cells were formed which lead to the formation of different tissues. The totipotency of the zygote is preserved into the germline cells while terminally differentiated cells can be found in the soma.

This important difference between CD- and CR-CLA perspective on cell differentiation also implies that, rather than repeatedly “self-renewing” developmental totipotency that has to be preserved upon repeated bifurcating cell fate differentiation divisions (Figure 6A), the developmental totipotency is intrinsically maintained in the ancestor zygote which ends into germline cells (Figure 6B).

## DISCUSSION

Since the discovery of repeated cell reproductions (CR) by a same mother cell (Figure S3-6) a CR-based cell lineage analysis (CLA) has been utilized for constructing genealogy-revealing cell linage for unicellular prokaryotic organisms (Liu, 1999) as well as for multicellular eukaryotic organisms (Liu, 2005b; and Liu, 2011a and b). The CR-CLAs for unicellular prokaryotic organisms were based on real-time continuous tracking of individual growth and reproduction of bacteria (Liu, 1999) (Figure S7). The CR-CLAs for multicellular eukaryotic organisms were first performed *in silico* with a generic theoretical model (Liu, 2005b) (Figure S8). Later a publication presenting a CD-CLA for zebrafish embryogenesis (Olive et al., 2010) was criticized (Liu, 2011a). In this criticism it was pointed out that the CD-based cell lineage reconstruction did not reflect true genealogy because it contained many “ghost mother” cells – the mother cells that should not exist by the cell “division” principle and were actually renamed into daughter cells. With the existence of these extra “ghost mother” cells, a CD-based cell lineage for the 64-cell stage zebrafish embryogenesis actually contained 127 cells (Figure S9). However, when the same original data was analyzed from a CR perspective, cell lineage was reconstructed with the exact same number of cells for each stage of zebrafish embryogenesis (Liu, 2011b) (Figure S10). Furthermore, a clear genealogy for zebrafish embryogenesis from zygote to the 64-cell stage was shown with a pedigree tree (Figure S11).

Unfortunately, the neglect by the mainstream on CR discovery and the resistance for publishing these insightful CR-CLAs in the established journals (Liu, 2012b) has resulted in a continued dominance of mistaken CD-CLA for cell lineage reconstructions. AceTree, a popular tool for visual analysis of *Ce* embryogenesis by linking images and annotations using tree representation, still follows the CD principle of one mother cell dividing into two daughter cells (Bao et al., 2006; Boyle et al., 2006; Murray and Bao, 2012). In order to force a cell lineage across a generation in the tree, one “sister” cell was renamed from a previous mother cell which was considered as “dead” by the cell division principle. For example, the left daughter from the division of Aba cell was tracked as ABal because the latter occupied the location of the former (Boyle et al. 2006). However this ironical change (in name) of a “dead” mother cell into a “newborn” daughter cell in AceTree still does not overcome CD-CLA’s intrinsic deficiency in reflecting true genealogy. This is because AceTree does not make M-D distinction between two cells derived from one cell. More significantly, the “dead” mother cell was most likely tracked into a true daughter cell. This possibility exists as the current study showed a systematic error of mistaken M-D assignment following the conventional “left mother, right daughter” rule (Figure 1, 3, 4).

CD dogma restricts any generation and cell age distinction between two cells derived from one cell. But CR discovery dictates an absolute generation and age asymmetry between two cells derived from one cell no matter how similar these two cells look alike (Liu, 1999). This cell generation and age asymmetry between two cells derived from one cell is very critical for considering cell (“division” / reproduction) synchronization methodology (Liu 2004a and 2005c), especially when the two cells derived from one cell show some time lag in their later “division” / reproduction (Liu, 2004b and c). The existence of a generation gap and a chronological age difference between any “division”-paired or reproduction-linked cells suggests that single-cell study is necessary (Liu, 2005d) and pooling material from the paired cells or averaging data on paired cells to report as representation or average of the “same type” of cells may be inappropriate. Unfortunately, many single-cell studies just performed their experiments and/or analyzed their data in this erroneous way (Liu, 2014a).

As pointed out many times in this study a correct M-D assignment is very critical for reliable CR-CLA. Chronological cell age determination thus would be essential for reliable CR-CLA. There are potentially many ways to determine cell age. A hypothesis that older template DNA strands are retained by mother cells and younger template DNA strands are segregated into daughter cells upon cell reproductions pointed to the utility of DNA age as an indicator for cell age (Liu, 1999). An *in silico* DNA-labeling experiment was even presented to show its utility (Liu, 2005a) (Figure S12). This hypothesis explained the observation of semi-conservative DNA replication (Meselson and Stahl, 1958) from a different perspective (Liu, 2006a) (Figure S13). More importantly, when published data claiming random segregation of DNA strands (Kiel et al., 2007; Steinhauser et al., 2012; and Yadlapalli et al., 2011) were analyzed from a CR perspective, consistent patterns of retaining older template DNA strands by mother cells and segregation of younger template DNA strands into daughter cells were observed (Liu 2007a and b; Liu, 2012b; Liu, 2014c) (Figure S14-16). Thus, future studies on *C. elegans* cell lineage should examine this DNA-Cell aging axis in *C. elegans* development and make reliable M-D assignment to reproduction-linked cell pairs in later embryonic stages to extend truthful genealogy revelation for full embryo and beyond.

Technology advancement has made cell lineage tracking on *Ce* embryogenesis very convenient and easy (Thomas, et al., 1996; Heid et al., 2002; Giurumescu and Chisholm, 2011; Verbrugghe and Chan, 2011; and Mace, 2013). But a paradigm change from CD-CLA to CR-CLA is necessary for realizing tremendous value embedded in massive information generated from extensive research on *C. elegans* (Stain et al., 2001; Bao et al., 2006; Boyle et al., 2006; Murray and Bao, 2012; Santella et al., 2016; Zacharias and Murray, 2016).

When the life of a microbial prokaryotic unicellular organism (Figure S17) and the life of a symbolic macrobial eukaryotic multicellular organism (Figure S18) are compared from a common cell reproduction perspective, it becomes clear that there should be no dichotomy in terms of their basic living principles. A universal life model (Figure S19) can be applied to all forms of cellular lives which abide some common living principles including: (1) having a beginning and an ending; (2) developing through three life stages: pre-reproduction juvenile stage, reproductive adult stage, and post-reproduction senescent stage; and (3) aging with increasing chronological age. The distinction between unicellular and multicellular organism resides in the absence (Liu, 2006c; Liu, 2014b) (Figure S17) and the presence (Figure S18) of cell differentiation within an individual body of respective organism.

The publications of CD-CLA on *C. elegans* (Sulston and Horvitz, 1977; Sulston et al., 1983) established a roadmap for systematic study of genetic mutations on multicellular development (Horvitz and Sulston, 1980; Sulston 2003) and making a little worm a giant model organism for life science research (Goldstein, 2016; Kemphues 2016). I wish the publication of this CR-CLA on the very beginning stage of *C. elegans* embryogenesis could result in a rapid death of the CD dogma and warm embracement for CR new biology (Liu, 2006b). A call for creating human-cell atlas was made very recently (Hayden, 2016). Could a fast accomplishment of creating worm-cell atlas with CR-CLA provide not only a solid confirmation for CR-based life science but also some valuable experiences for human-cell atlas project?

## MATERIALS AND METHODS

CD-CLAs performed in this study were based on published information retrieved from respective research papers cited in the relevant Result sections. CR-CLAs of the current study used the same data from the respective CD-CLAs but interpreted those data in the CR ways described in earlier publications [Liu 1999, 2005a]. Briefly, the naming started with a single letter Z for zygote and added one alphabetical letter to a daughter cell following the alphabetical order. For example, the first and the second daughter of Z were named as Za, and Zb, respectively. This pattern of cell naming was continued for all later descendants. Thus, the number of letter(s) contained in a cell name directly reflects the generation of the cell. A superscript number in cell name indicates the number of reproduction experienced by the cell.

## ACKNOWLEDGEMENTS

I thank my wife Jing Zhang for decades-long support to my off-duty spare-time research on cell reproduction-based life science which was not funded by any grant. I also thank Chang-Zucherburg Initiative for setting the creation of “human-cell atlas” as its focus (https://www.facebook.com/chanzuckerberginititive/) which encouraged me to wrap up my decades-long amateur research on *Ce* even though it may still be regarded as “premature”.

This paper is dedicated to my parent and my children especially my father recently deceased and my son still struggling against multiple developmental abnormalities since his birth 26 years ago.

